# Relationship Between Motor Capacity of the Contralesional and Ipsilesional Hand Depends on the Side of Stroke

**DOI:** 10.1101/635136

**Authors:** Rini Varghese, Carolee J. Winstein

**Affiliations:** Division of Biokinesiology and Physical Therapy, University of Southern California; Department of Neurology, Keck School of Medicine, University of Southern California

**Keywords:** Ipsilesional deficits, stroke, hemispheric differences

## Abstract

There is considerable evidence that after a stroke, ipsilesional deficits increase as contralesional impairment increases. Here, we asked if the relationship between the motor capacities of the two limbs differs based on the side of stroke. Forty-two pre-morbidly right-handed chronic stroke survivors (left hemisphere damage, LHD = 21) with mild-to-moderate paresis performed distal items of the Wolf Motor Function Test (dWMFT). We found that compared to RHD, the relationship between contralesional arm impairment (Upper Extremity Fugl-Meyer, UEFM) and ipsilesional hand motor capacity was stronger (*R*^2^_*LHD*_= 0.42; *R*^2^_*RHD*_ < 0.01; *z* = 2.12; *p* = 0.03) and the slope was steeper (*t* = −2.03; *p* = 0.04) in LHD. Similarly, the relationship between contralesional dWMFT and ipsilesional hand motor capacity was stronger (*R*^2^_*LHD*_= 0.65; *R*^2^_*RHD*_= 0.09; *z* = 2.45; *p* = 0.01) and the slope was steeper (*t* = 2.03; *p* = 0.04) in LHD compared to RHD. Multiple regression analysis confirmed the presence of an interaction between contralesional UEFM and side of stroke (*β*_3_= 0.66 ± 0.30; *p* = 0.024) but only trended towards significance for the interaction between contralesional dWMFT and side of stroke (*β*_3_= −0.51 ± 0.34; *p* = 0.05). Results were confirmed after removal of potential outliers. Our findings suggest that the relationship between contra- and ipsi-lesional motor capacity depends on the side of stroke, such that the inter-limb relationship is stronger for stroke survivors with left hemisphere damage compared to those with right hemisphere damage.

## Introduction

It is now well known that unilateral stroke not only results in contralesional arm deficits, but also significant albeit more subtle motor deficits in the ipsilesional limb compared to age-matched non-disabled adults.^1–4^ Previous work that examined the relationship between motor capability or capacity of the contralesional and ipsilesional hands are in agreement that the presence of motor deficits in the ipsilesional arm and hand are related to the severity of motor deficits in the contralesional upper limb, especially in the chronic phase after stroke.^5–9^ For example, Boyd and colleagues demonstrated that the ability to learn an implicit motor task with the ipsilesional hand was inversely correlated with the degree of motor impairment in the contralesional upper extremity.^5^ Rinne and colleagues^6^ used a force-tracking task and showed that grip strength and tracking accuracy co-varied between the ipsilesional and contralesional hands.^6^ Similarly, two recent studies reported that deficits in the ipsilesional hand were most impaired for a group classified with relatively more “severe” contralesional motor impairments, defined as an Upper Extremity Fugl-Meyer score (UEFM) of less than 28,^7^ or moderate impairment, 32 to 57.^10^

In addition to the studies aimed at characterizing the relationship between contralesional and ipsilesional hands, there is mounting evidence that the unilateral motor deficits observed for contralesional and ipsilesional limbs are hemisphere-specific and thus depend on side of stroke lesion.^8,11–16^ For example, using clinical motor assessments of grip strength and hand dexterity, Harris and Eng^13^ showed that in chronic stroke survivors who are pre-morbidly right-hand dominant, contralesional motor impairments were less severe in individuals who suffered damage in the dominant (i.e. left) hemisphere (LHD) compared to those who suffered damage in the non-dominant (right) hemisphere (RHD).^13,16^ In contrast, considering ipsilesional motor deficits, the evidence is mixed concerning hemisphere-specific effects. For instance, some studies reported that individuals with LHD exhibited more severe ipsilesional arm and hand deficits compared to those with RHD ^4,16–18^ while others have reported no difference in ipsilesional capacity between LHD and RHD.^2^ In acute stroke survivors, Kust et al demonstrated that deficits in grip force of the ipsilesional hand were significantly associated with clinical measures of function of the contralesional hand *only* in LHD.^14^ Contrary to this, de Paiva Silva et al found that the ipsilesional hand was significantly slower and less smooth in individuals with RHD who exhibited moderate-to-severe motor impairments (UEFM < 34) in the contralesional upper extremity compared to controls, LHD, and those with mild motor impairment.^8^

Taken together, there is converging evidence regarding the relationship between motor deficits of the contralesional and ipsilesional upper extremity, such that ipsilesional deficits are worse when contralesional impairment is greater (Figure 1A); however, it is uncertain whether this relationship between the two limbs depends on which hemisphere is damaged. In particular, motor deficits of the two limbs are most prominent for tasks that require dexterous motor control (e. g., grip force, tapping, tracking). For predominantly right-handed cohorts (as is the case in most studies), contralesional deficits appear to be more severe in those with RHD, in whom the contralesional limb is non-dominant; whereas ipsilesional deficits were more severe in those with LHD. An exception to this observation for those with RHD seems to be in the case when contralesional impairment is most severe (i.e., UEFM < 34).^7,8^ Thus, one might predict that as contralesional impairment worsens, individuals with LHD would have proportionally worse ipsilesional deficits, but individuals with RHD (especially if say UEFM > 34) would not; see (Figure 1 B & C) for two alternative hypotheses. This prediction arising from an interaction between severity of contralesional deficits and the hemisphere affected by the stroke has not before been explicitly tested.

**Figure 1.**
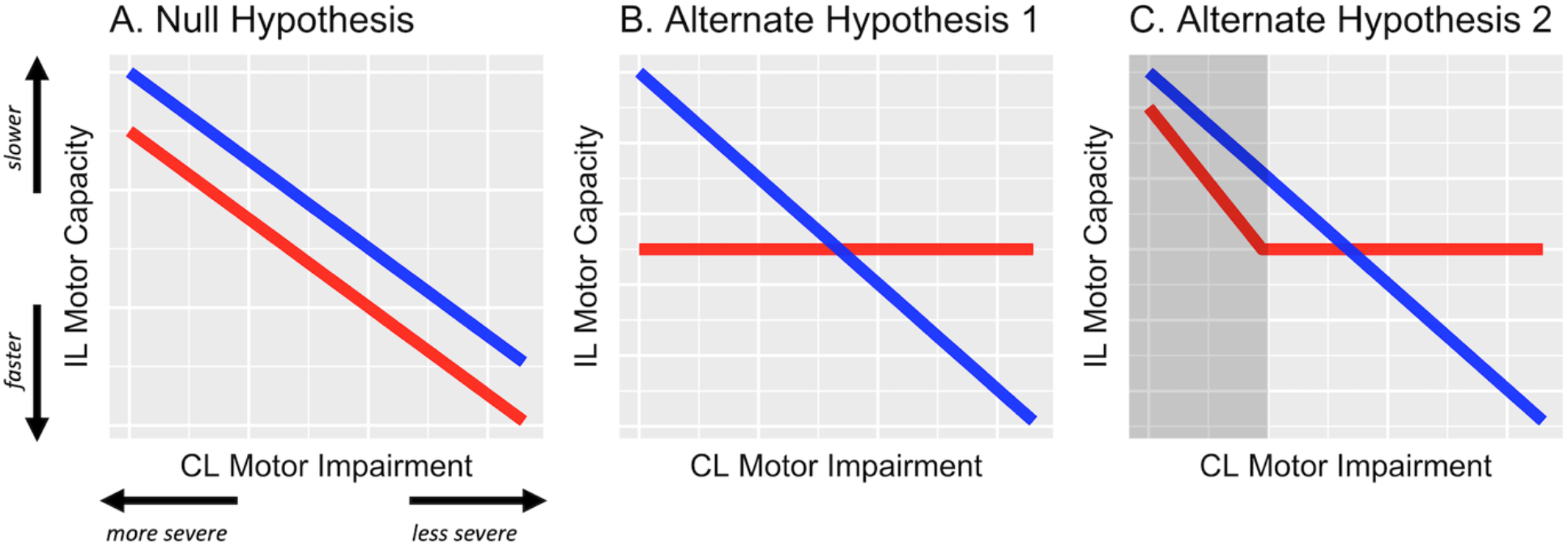
Hypothesized effects represented in schematic figure. A. The null hypothesis, wherein the relationship between contralesional (CL) impairment and ipsilesional (IL) motor capacity is not modified by the side of stroke lesion. B. Alternative hypothesis 1, wherein ipsilesional deficits are related to contralesional impairment but only in LHD (blue) and not in RHD (red). C. Alternate hypothesis 2, wherein ipsilesional deficits are related to contralesional impairment but only in LHD and in RHD with severe impairment (represented in the shaded dark-grey area).

One reason that this prediction remains untested might be methodological in that in at least three of the studies discussed earlier, participants were categorically classified based on the degree of contralesional motor impairment (e.g., mild, moderate, severe).^7,8,10^ While grouping in this manner may be somewhat useful for stratification and randomization purposes with large samples, categorization (or worse, dichotomization) of a continuous variable around arbitrarily set cut-off points presents several concerns. Of concern is a loss of measurement resolution, an assumption of discontinuity in the underlying construct (in this case motor impairment), unequal subgroup sizes (or biased sampling), and large unexplained residuals in regression models, to name a few.^19–21^ Overall, if the objective is to understand the nature and extent of critical response-predictor relationships, then a categorical approach is particularly problematic.

Thus, the primary objective is to determine if the severity of deficits in the ipsilesional hand varies directly with that of the contralesional hand (using a continuous measure). Further, and more importantly, we seek to determine if this relationship differs based on the side of stroke lesion (i.e., an interaction effect). We predict that motor capacity of the ipsilesional hand will vary directly with the severity of the contralesional motor impairment and dexterous motor capacity, only in individuals with LHD, but not in individuals with RHD; see (Figure 1).

## Methods

### Participants

Forty-two chronic stroke survivors (n = 21 left-hemisphere damage, LHD) provided informed consent to participate and in accordance with the 1964 Declaration of Helsinki and the guidelines of the Institutional Review Board for the Health Science Campus of the University of Southern California. These participants were enrolled as part of a larger phase-IIb clinical trial (Dose Optimization for Stroke Evaluation, ClinicalTrials.gov ID: NCT01749358).

Participants were included if they were: 1) between 21-75 years of age, 2) pre-morbidly right-handed, 3) ≥ 150 days post stroke (chronic phase), 4) had mild to moderate residual motor impairment (Upper Extremity Fugl-Meyer, UEFM score ≥ 19) with mostly resolved upper extremity paresis. Participants were excluded if they had: 1) severe sensory disturbances (no response to light touch or complete loss of proprioception as indicated by the UEFM), 2) current major depressive disorder (score > 3 on PHQ2, depression screening survey) 3) a history of recent surgeries, significant orthopedic injuries, or pain affecting the upper extremity that would restrict shoulder and elbow movement, 4) severe cognitive deficits such as aphasia, apraxia or neglect that would preclude participants from comprehending test instructions or questionnaires.

### Outcome Measures

#### Motor Component of the Upper Extremity Fugl-Meyer (UEFM)

The UEFM ^22^ is an assessment of motor impairment of the contralesional arm and hand after stroke and includes tests of strength and independent joint control. Item-wise scoring of the UEFM ranges from 0 (unable to perform) to 2 (able to perform completely) while total score ranges from 0 to 66, with a higher score indicating lesser impairment.

#### Wolf Motor Function Test (WMFT)

The WMFT is designed to assess upper extremity motor capacity through timed functional task performance (e.g., lifting a can, pencil, or paper clip). Originally designed for patients with moderate to severe upper extremity motor deficits, the test was later modified by Morris, Crago and Taub to accommodate individuals with mild motor impairments ^23^. In a series of 15 tasks, the test administrator asks the participant to perform frontal or midsagittal plane motions with the shoulder and elbow, and dexterous tasks with the hand. Item-wise scoring entails a continuous capacity time-score. Traditionally, the test has been used to assess motor capacity of both the contralesional as well as the ipsilesional arm and hand, with the latter used as a reference for comparison within an individual.

We used the WMFT time-score to assess upper extremity motor capacity through timed functional task performance. A principle component analysis of WMFT scores revealed two clusters: one consisting of the proximal (#1-8, except 6, i.e., lifting weight to box), the other consisting of the distal (#9-17, except 14, i.e., grip strength),^24^ with the latter serving as the primary measure of hand motor capacity. The distal battery (dWMFT) consists of the following 8 tasks: lift can, lift pencil, lift paper clip, stack checkers, flip cards, turn a key in a lock, fold towel, and lift basket. Hand motor capacity was assessed in both limbs.

### Statistical Analyses

All analyses were conducted in the R statistical computing package version 3.5.1.^25^ To test the hypothesis that the inter-limb relationship of motor capacity is modified by the side of stroke lesion, we used the coefficient of determination (*R*^2^) and compared the covariances between LHD and RHD using the Fisher’s Z test. We then performed a simple linear regression to determine the slopes of the relationship between contralesional (CL) UEFM and ipsilesional (IL) dWMFT (Model 1), and, CL dWMFT and IL dWMFT (Model 2). We used t-tests to compare these slopes between LHD and RHD.

To supplement these primary analyses and as a more robust assessment of the interaction between the side of lesion and contralesional motor capacity, we used multiple linear regression of the following form:

**Model 1**: *y* = *β*_0_ + *β*_1_(*UEFM*) + *β*_2_(*side of stroke*) + *β*_3_(CL *UEFM* ∗ *side of stroke*) + *ϵ*
**Model 2**: *y* = *β*_0_ + *β*_1_(*CL dWMFT*) + *β*_2_(*side of stroke*) + *β*_3_(CL *dWMFT* ∗ *side of stroke*) + *ϵ*

In both models, *y* is the average time score on the distal WMFT of the ipsilesional hand. Using this multiple model, our hypotheses were that *β*_1_ ≠ 0 and *β*_3_ ≠ 0 (see Figure 1). Any statistically significant interaction was resolved post-hoc using a t-test comparison of estimated marginal trends between LHD and RHD.

All continuous variables were assessed for normality using Lilliefors test (modified Kolmogorov-Smirnov test). Distributions for chronicity and average time-score for the distal WMFT were negatively skewed and were therefore log-transformed. Welch’s t-tests were used to compare age, chronicity, and Upper Extremity Fugl-Meyer scores between LHD and RHD, whereas Chi-square test was used to compare the proportion of females and males between the two groups. Each group was standardized to its own unit variance (z-scored) to equalize range and for subsequent linear regression analysis. Outliers were identified by visual inspection of scatterplots. Any value of IL dWMFT more extreme than ± 1.5 log-SD was examined carefully for their influence on interlimb covariance and slopes. If removal of these observations did not change the direction or significance of the effect in the simple model, we included them in the final model. Residuals of the final model were analyzed to confirm that all necessary assumptions for multiple regression were met. Significance level (α) was set at *p* = 0.05.

In order to select the predictor variables that best explain the response, we used a backward selection approach, in which we began by adding all predictor variables in each of the two above models to explain the response variable *y*. This included our hypothesized predictors, CL UEFM (or CL dWMFT) and the side of stroke lesion (LHD or RHD), and, potential confounders (age, chronicity, and sex). In a combined full model, those predictors that met a liberal cut-off of *p* = 0.2 were preserved in the final reduced model. Based on this selection process, we found sex to be a significant confounder (*p* = 0.08) in Model 1, and therefore included it as a predictor in the reduced Model 1. For Model 2, none of the confounders met the cut-off p-value, except our hypothesized predictors. Additional information on model selection and model diagnostics is included as supplementary materials. Standard errors and 95% CI of the estimates of regression coefficients were confirmed by performing 1000 bootstrap replicates.

## Results

Descriptive statistics for all participants are provided in Table 1. On average, the 42 stroke survivors had moderate arm impairment (UEFM = 41.6), were approximately 60 years of age, 5.75 years post-stroke, and were predominantly male (74%). There were no significant differences between LHD and RHD in the level of impairment, chronicity or the number of males. Individuals with RHD were younger compared to LHD (median age difference 8.7 years) but not statistically different.

**Table 1.** Descriptive Statistics for the full sample (N = 42), and for the two groups of interest, left hemisphere damage, LHD (n = 21) and right hemisphere damage, RHD (n = 21).

| Variable | Overall (N = 42) | LHD (n = 21) | RHD (n = 21) |
| --- | --- | --- | --- |
| | Mean * (± SD) | Mean (± SD) | Mean (± SD) |
| Sex (Male) * | 31 (73.8) | 15 (71.4) | 16 (76.2) |
| Age (in years) | 59.16 (12.3) | 60.71 (10.8) | 57.62 (13.8) |
| Chronicity (in months) | 69.37 (36.9) | 63.32 (24.4) | 75.41 (46.1) |
| UEFM Motor (/66) | 41.57 (10.5) | 40.67 (12.6) | 42.48 (8.2) |

\* Count (percent) for categorical variables

| **Variable** | **Overall (N = 42)**<br>Mean * (± SD) | **LHD (n = 21)**<br>Mean (± SD) | **RHD (n = 21)**<br>Mean (± SD) |
| --- | --- | --- | --- |
| Sex (Male) * | 31 (73.8) | 15 (71.4) | 16 (76.2) |
| Age (in years) | 59.16 (12.3) | 60.71 (10.8) | 57.62 (13.8) |
| Chronicity (in months) | 69.37 (36.9) | 63.32 (24.4) | 75.41 (46.1) |
| UEFM Motor (/66) | 41.57 (10.5) | 40.67 (12.6) | 42.48 (8.2) |

\* Count (percent) for categorical variables

### Model 1: Side of lesion modifies the relationship between CL UEFM and IL motor capacity

Contralesional UEFM explained 42% of the variance in ipsilesional hand motor capacity in LHD (*p* = 0.001), but less than 1% in RHD (*p* > 0.05). The slope of this relationship was −0.65 ± 0.17 (*p* = 0.001) in LHD and −0.066 ± 0.23 (*p* = 0.78) in RHD. Compared to RHD, the covariance between contralesional UEFM and ipsilesional hand motor capacity was significantly stronger (Fisher’s *z* = 2.12, *p* = 0.03) and the slope was steeper in LHD (*t* = −2.03, *p* = 0.04).

Four observations (two each in LHD and RHD) were identified as potential outliers. After removal of these outliers, contralesional UEFM explained 44.3% of the variance in ipsilesional hand motor capacity in LHD (*p* = 0.001), and 2.26% in RHD (*p* > 0.05). The slope of this relationship changed to −0.42 ± 0.11 (*p* = 0.001) in LHD and 0.13 ± 0.20 (*p* = 0.54) in RHD. Again, a comparison of the covariances and slopes between the groups revealed that compared to RHD, the relationship between contralesional UEFM and ipsilesional hand motor capacity was significantly stronger (Fisher’s *z* = 2.7, *p* = 0.006) and the slope was steeper in LHD (*t* = −2.41, *p* = 0.02).

Since these observations did not significantly change the strength of covariance nor the slope of the relationship, they were preserved in the final multiple model. Analysis of residuals of the final model did not indicate violations of necessary assumptions in multiple regression in terms of linearity, equality of variance, independence and normality of errors, and multicollinearity of independent variables, nor the presence of unduly influential observations. Nonetheless, estimates below are reported both with and without suspected outliers.

After adjusting for main effects and significant confounders using multiple regression, the final reduced form of Model 1 was statistically different from a null model (*F* = 3.47, *p* = 0.016, adjusted *R*^2^= 0.19). Based on estimates from Model 1, CL impairment (UEFM) was significantly associated with IL hand motor capacity, i.e., dWMFT, (*β*_1_ = −0.72 ± 0.21, *p* = 0.001; *without outliers*: −0.44 ± 0.17, *p* = 0.01) (Figure 2A). There was no significant effect of the side of lesion (*β*_2_ = 0.026 ± 0.27, *p* = 0.92; *without outliers*: 0.22 ± 0.23, *p* = 0.33). There was a significant interaction between the side of lesion and CL impairment (*β*_3_ = 0.66 ± 0.30, *p* = 0.024; *without outliers*: 0.56 ± 0.24, *p* = 0.024). Post-hoc contrasts of estimated marginal trends indicated that the slope of the relationship between CL UEFM and IL dWMFT was significantly more negative in LHD compared to RHD (*t* = −2.34, p = 0.02; *without outliers*: −2.37, p = 0.02). Figure 2 B and C illustrates the interaction.

**Figure 2.**
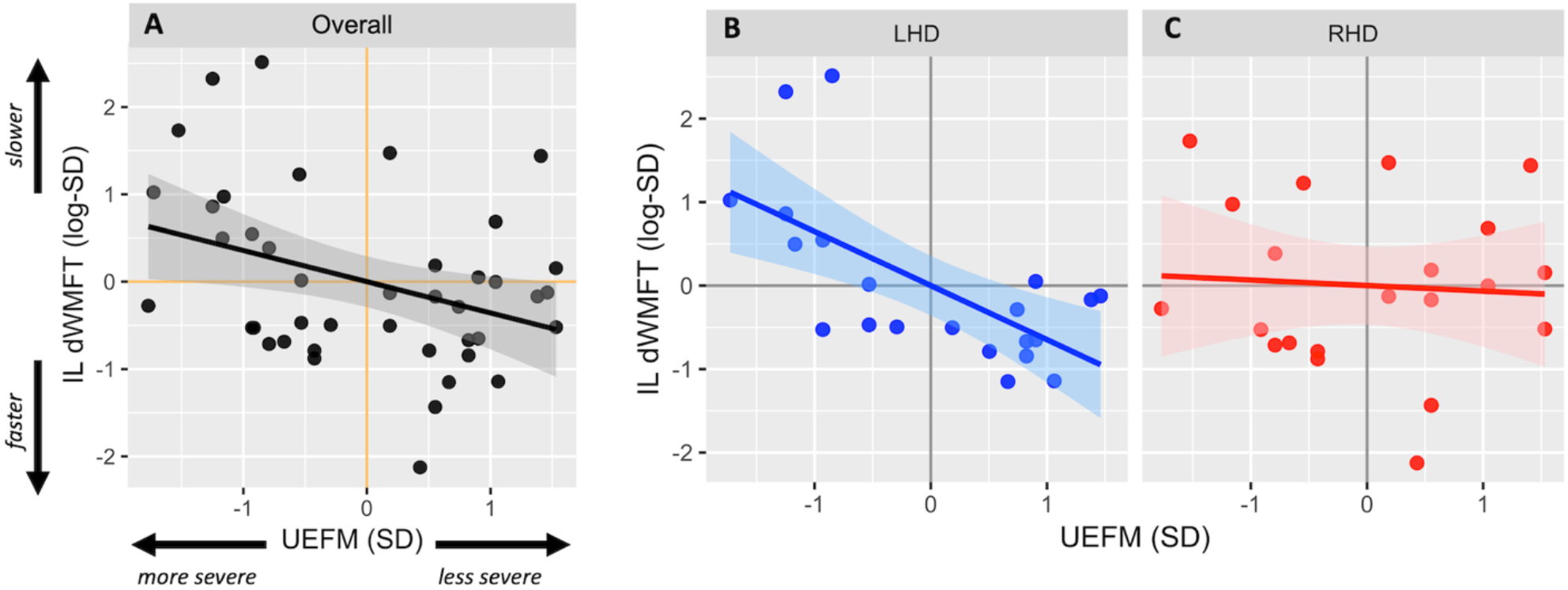
Scatterplots show the relationship between contralesional motor impairment (CL UEFM) and ipsilesional distal motor performance (IL dWMFT) for the full sample (A), LHD (B), and RHD (C). Solid lines represent the linear prediction and shaded areas represent the 95% confidence interval.

### Model 2: Side of lesion modifies the relationship between CL dWMFT and IL motor capacity

Contralesional dWMFT explained 65% of the variance in ipsilesional hand motor capacity in LHD (*p* < 0.001), but only 9% in RHD (*p* > 0.05). The slope of this relationship was 0.81 ± 0.13 (*p* < 0.001) in LHD and 0.29 ± 0.22 (*p* = 0.19) in RHD. A comparison of the covariances and slopes between LHD and RHD revealed that compared to RHD, the relationship between CL dWMFT and IL motor capacity was significantly stronger (Fisher’s *z* = 2.45, *p* = 0.01) and the slope was steeper in LHD (*t* = 2.03, *p* = 0.04).

After removing the outlying observations, contralesional dWMFT explained 62% of the variance in ipsilesional hand motor capacity in LHD (*p* < 0.001), and < 1% in RHD (*p* > 0.05). The slope of this relationship changed to 0.54 ± 0.1 (*p* < 0.001) in LHD and 0.05 ± 0.21 (*p* = 0.81) in RHD. Compared to RHD, the relationship between CL dWMFT and IL motor capacity was significantly stronger (Fisher’s *z* = 2.85, *p* = 0.004) and the slope was steeper in LHD (*t* = 2.11, *p* = 0.04).

Since these observations did not significantly change the strength of covariance or the slope of the relationship, they were preserved in the final multiple model. Once again, analysis of residuals did not indicate violations of necessary assumptions in multiple regression nor the presence of unduly influential observations. Nonetheless, estimates below are reported both with and without suspected outliers.

After adjusting for main effects and significant confounders using multiple regression, the final reduced form of Model 2 was statistically different from a null model (*F* = 7.48, *p* < 0.001, adjusted *R*^2^ = 0.32). Based on estimates from Model 2, CL hand motor capacity was significantly associated with IL hand motor capacity (*β*_1_= −0.81± 0.16, *p* < 0.001; *without outliers*: 0.54 ± 0.17, *p* = 0.003) (Figure 3A). There was no significant effect of the side of lesion (*β*_2_= 0.12 ± 0.26, *p* = 0.66; *without outliers*: 0.19 ± 0.22, *p* = 0.39). There was an interaction between the side of lesion and CL hand motor capacity, but it only approached statistical significance (*β*_3_ = −0.51 ± 0.34, *p* = 0.05; *without outliers*: −0.49 ± 0.24, *p* = 0.046). Figure 3 B and C illustrates the interaction.

**Figure 3.**
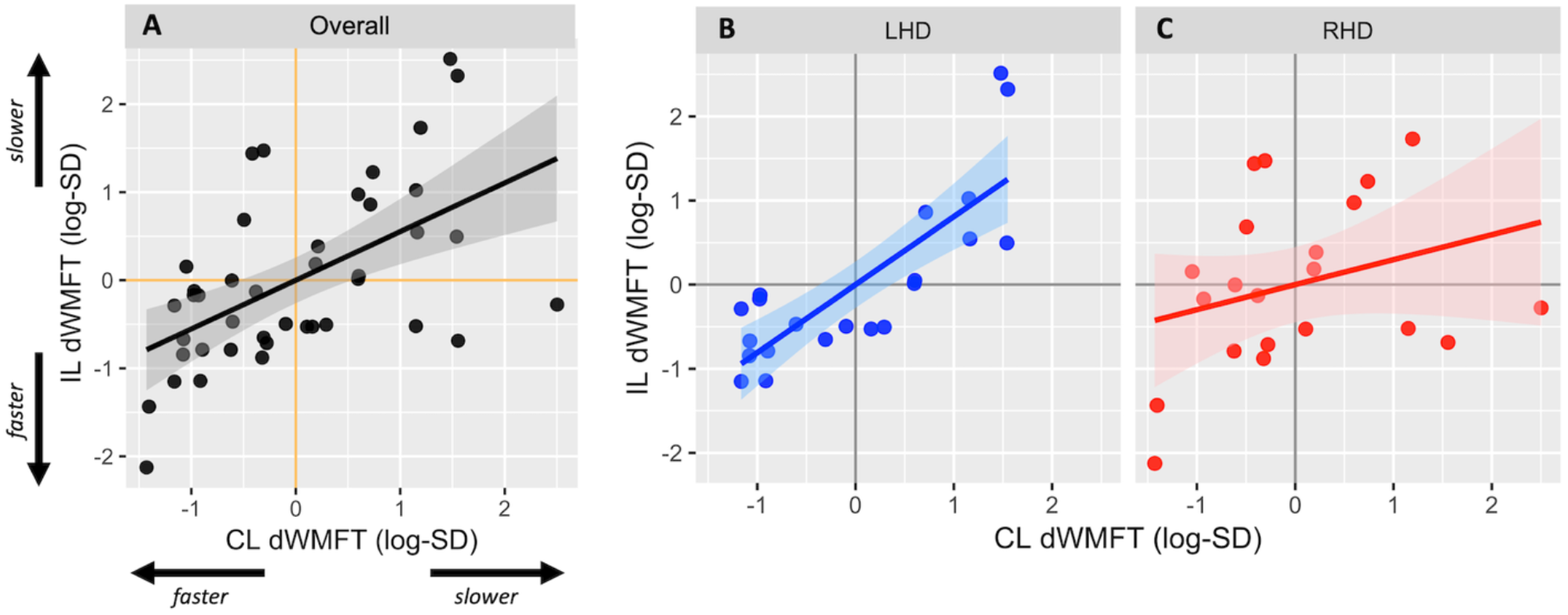
Scatterplots show relationship between contralesional distal motor performance (CL dWMFT) and ipsilesional distal motor performance (IL dWMFT) for the full sample (A), LHD (B), and RHD (C). Solid lines represent the linear prediction and shaded areas represent the 95% confidence interval.

## Discussion

For the first time, we explicitly tested the hypothesis that motor capacity of the ipsilesional hand is influenced by an interaction between the severity of contralesional deficits and the side of stroke lesion. We found that ipsilesional motor capacity co-varies with contralesional impairment to a significantly greater degree in individuals with LHD compared to RHD. Hints of this interaction were implicit in a few previous studies;^2,3,15^ however, categorical reporting of the Upper Extremity Fugl-Meyer (UEFM) masked this interesting effect. In the following sections, we provide an analysis of the interaction effect, explore the insights obtained from the type of task evaluated, and discuss the role of the left hemisphere in the organization of motor outputs for both limbs. Finally, we identify the limitations of this study and suggest future research questions.

### Analysis of the interaction effect

By preserving the continuity of the UEFM and utilizing continuous standardized z-scores, our statistical approach allowed a direct comparison of our regression parameters (i.e., *βs*), thus reflecting effect sizes of each of the candidate predictors—contralesional impairment, side of lesion and their interaction. While contralesional impairment alone bore the largest effect on ipsilesional motor capacity (*β*_1_), the second largest effect was through the interaction between contralesional UEFM and the side of lesion (*β*_2_), and not the side of lesion alone (*β*_3_). In fact, unlike previous findings, we did not observe a significant effect of the side of lesion on ipsilesional motor capacity ^17,18^, nor on contralesional UEFM or dWMFT.^13^

One reason for this might be that the effect of the side of lesion observed in those previous studies may have arisen from its interaction with contralesional impairment. However, because an interaction effect was not explicitly tested and because contralesional impairment was either collapsed across the groups ^2,3,15^ or categorical,^7,8^ variance in the ipsilesional capacity may have been conflated with the effect of the side of lesion, or, remained unexplained, especially for UEFM scores that fell at the boundaries of the pre-defined categories. To illustrate this point further, let us revisit Figure 1B. Here, if a cut-off for the x value (say, UEFM) was to be set at any point to the left of where the two lines intersect, one might conclude that ipsilesional deficits are greater in LHD compared to RHD, whereas a cut-off to the right of the intersection might suggest that ipsilesional deficits are greater in RHD. Both observations would indicate an effect of the side of lesion, but not the change in the direction of this effect, evident as an interaction.

Naturally, this brings to light another important issue and that is the choice of cut-off scores for categorization, as this would be crucial in determining the direction of the effect of the side of lesion. Previously, various cut-off scores for the UEFM have been used to define impairment categories.^7,8,10^ Of these, our data suggest that a UEFM score of 42, which occurs at the intersection of the linear fits for LHD and RHD, would best reflect the change in the direction of effect, i.e., the interaction between side of lesion and contralesional impairment on ipsilesional capacity. Interestingly, *at* this score of 42, there would appear to be no differences in motor capacity of either hand between LHD and RHD, which might explain why a number of large clinical trials (e.g. EXCITE,^26^ ICARE ^27^), designed for mild to moderately impaired stroke survivors (mean UEFM scores in these studies were 42.5 and 41.6 respectively) may not have observed, on average, any differences in motor capacity based on the side of stroke.

### Insights from the type of task

A common link between our study and past reports is that ipsilesional deficits were found to be most pronounced for distal (dexterous) tasks. These tasks—lift can, lift pencil, lift paper clip, stack checkers, flip cards, turn a key in a lock, fold towel, and lift basket—nearly always involve object manipulation and inherently require dexterous motor control of the hands. Sunderland and colleagues^4^ demonstrated that early on after a stroke, spatial accuracy in dexterity tasks performed with the ipsilesional hand correlated with cognitive deficits, such as apraxia, in individuals with LHD. While individuals included in this study did not exhibit severe apraxia and were approximately 5 years post-stroke, it is possible that mild cognitive deficits, including apraxia, may have impacted dexterous task performance in those with LHD, especially in the more severe ranges of UEFM. Furthermore, we note that our evaluation of dexterous task performance was through timed tests, and not quality of movement or accuracy. It has been suggested that the left hemisphere plays an important role in regulating the timing and speed of movements^15^, and thus, injury to the left hemisphere, particularly to premotor and fronto-parietal networks (e.g. IPC)^27, 28^ may impair planning and sequencing required for smooth and rapid performance of dexterous motor tasks.

That deficits are apparent for distal rather than proximal motor performance is also relevant to our understanding of the putative neural substrates responsible for ipsilesional deficits. In this regard, candidate substrates include direct ipsilateral corticofugal pathways or indirect inhibitory circuits in the cortex, sub-cortex (via the corpus callosum and other commissures), or spinal cord (via spinal inter-neuronal circuits). As for the indirect cortical inhibitory circuits, the classic view contends that, based purely on somatotopy, the control of distal musculature, represented laterally on the motor cortex, lacks transcallosal fibers. Thus, control of the ipsilesional hand (or lack thereof) might be attributable to intra-hemispheric pathways rather than interhemispheric interactions. Whereas this view has been challenged based on empirical neurophysiologic evidence in animals and humans,^30,31^ the presence of ipsilateral activations in acallosal patients^32^ provides further reason to suspect that intra-hemispheric pathways, or perhaps inhibitory circuits in the lower centers of the neuraxis, are likely responsible for the control of distal musculature of the ipsilateral hand. For a comprehensive review of this topic, please see Carson (2005).^33^ Nevertheless, whether this type of ipsilateral control is normal or maladaptive is a controversial topic^34–36^ and warrants further study.

### The role of the left hemisphere in the control of both hands

Our main observation that deficits in ipsilesional hand motor capacity scale with contralesional impairment only in LHD is qualitatively similar to previous clinico-behavioral ^e.g. 17,18,37,38^ and phenomenological evidence. ^e.g. 39–41^ These findings are consistent with a rather simplified organizational model of the nervous system in which certain aspects of motor and/or cognitive control are lateralized to the left (or dominant) hemisphere, such that damage to the left hemisphere results in deficits in skilled motor actions of both upper extremities. For example, using EMG recordings of homologous muscles in the arm, Cernacek (1961) demonstrated that the frequency of motor irradiations, i.e., unintended motor output in the ipsilateral hand, were significantly higher from the dominant to the non-dominant extremity.^39^ Similarly, Wyke (1968) reported that while individuals with left-sided cerebral lesions exhibited bilateral motor deficits in speed and limb postural control, deficits in those with right cerebral lesions were restricted to the contralateral limb.^18^ Lastly, in one of the earliest experiments using functional MRI, Kim and colleagues (1993) showed that the task-evoked activation of the left hemisphere was substantially greater for ipsilateral movements compared to the right hemisphere.^42^ In later years, a number of neuroimaging ^43,44^ and neurophysiologic^45–47^ studies have provided confirmatory evidence for the role of the dominant hemisphere in organizing motor outputs to both hands. Our results of co-varying deficits between the contralesional and ipsilesional hand in LHD provides further empirical support for the role of the left hemisphere (in our pre-morbidly right-handed group) in the control of both hands.

### Limitations and future considerations

Some of the most significant methodological shortcomings of this study are that: First, the study design was purely observational, and the statistical analyses were inferential with a relatively small sample size. To this point, we conducted analyses with and without outliers, and found that while exclusion of outliers affected the strength of the overall model, it did not affect the probability associated with rejecting the null hypothesis. A prospective study or independent validation in a separate cohort would be ideal, if larger samples were available. A larger sample would render more robust findings that are less sensitive to distortions from outlying values.

Second, UEFM scores for the RHD group were restricted towards the more severe range, with the most severely impaired individual’s score being 28. This restriction, however, was less so in the LHD group (min. UEFM = 19). Although this limitation in range was circumvented by using group-wise z-scores, we are cautious in generalizing our observations regarding the interaction effect to more severe ranges of motor impairment in RHD. This is quite apparent in the variability around our estimated linear fits especially towards the extreme ranges of predictor values for RHD. Indeed, it is possible that for the more severe range in RHD, there exists a linear relationship between contralesional and ipsilesional motor deficits as illustrated in Figure 1C. Thus, while we can, with some confidence, reject the null hypothesis (Figure 1A), our data are insufficient to differentiate between the two alternate hypotheses, and warrant a follow-up study.

It must be emphasized that the absence of a relationship with contralesional impairment in RHD should not be taken to mean that ipsilesional deficits are absent in this group. In fact, there is substantial evidence to the contrary. Comparison with an appropriate control group would be necessary to demonstrate the presence of ipsilesional deficits in RHD and the functional implications of these deficits. As alluded to earlier, measuring the speed of performance, as in the case of timed functional tasks assessed here, does not provide specific information about perceptual errors, spatial accuracy or visuomotor deficits, which, based on previous evidence,^48,49^ might be a more important indicator of motor performance in RHD.

In summary, our results suggest that ipsilesional motor deficits co-vary with the degree of impairment in LHD but is less pronounced in RHD. This observation further underscores the extensive motor experiences of the pre-morbidly dominant ipsilesional limb and the importance of the left hemisphere in the control of timed tasks for both hands. For the future, we propose that a hypothetical model of bilateral deficits in LHD is readily testable through a prospective study that uses a bimanual experimental paradigm with sensitive kinematic measures. Such a paradigm could offer important insights into the role and organization of each hemisphere for the control of uni- and bi-manual movements.

## Supplementary Material

The complete data table and codebook for statistical analysis, including additional resources, is available with this manuscript as well as through the first author’s OSF Repository: https://osf.io/pbtk9

